# Multifractal signatures of perceptual processing on anatomical sleeves of the human body

**DOI:** 10.1101/2020.05.12.091702

**Authors:** Madhur Mangalam, Nicole S. Carver, Damian G. Kelty-Stephen

## Abstract

Research into haptic perception typically concentrates on mechanoreceptors and their supporting neuronal processes. This focus risks ignoring crucial aspects of active perception. For instance, bodily movements influence the information available to mechanoreceptors, entailing that movement facilitates haptic perception. Effortful manual wielding of an object prompts feedback loops at multiple spatiotemporal scales, rippling outwards from the wielding hand to the feet, maintaining an upright posture, and interweaving to produce a nonlinear web of fluctuations throughout the body. Here, we investigated whether and how this bodywide nonlinearity engenders a flow of multifractal fluctuations that could support perception of object properties via dynamic touch. Blindfolded participants manually wielded weighted dowels and reported judgments of heaviness and length. Mechanical fluctuations on the anatomical sleeves, from hand to the upper body, as well as to the postural center of pressure, showed evidence of multifractality arising from nonlinear temporal correlations across scales. The modeling of impulse-response functions obtained from vector autoregressive (VAR) analysis revealed that distinct sets of pairwise exchanges of multifractal fluctuations entailed accuracy in heaviness and length judgments. These results suggest that the accuracy of perception via dynamic touch hinges on specific flowing patterns of multifractal fluctuations that people wear on their anatomical sleeves.

## 1. Introduction

### 1.1 Modulating the bodywide flow of mechanical fluctuations to investigate haptic perceptual performance

Research into haptic perception typically concentrates on mechanoreceptors and their supporting neuronal processes, such as mechanoreceptor physiology and neuronal processing of passive somatosensory feedback [1,2]. Despite the significant insights of this research, this focus risks ignoring crucial aspects of active perception. For instance, bodily movements influence the information available to mechanoreceptors, entailing that movement facilitates haptic perception [3–5]. The present work investigated how bodywide mechanical interactions facilitate “dynamic” or “effortful” perception of heaviness and length of manually-wielded, visually-occluded objects. Specifically, we test two possibilities: first, that statistical structure in mechanical fluctuations flows across disparate anatomical locations (i.e., beyond the wielding hand) to coordinate perceptual judgments and, second, that the structure of this flow of statistical regularities impacts the accuracy of these judgments.

### 1.2 The bodywide multifractal tensegrity (MFT) may simplify the degrees-of-freedom problem of spatiotemporally organizing afferent activity

The human body is highly complex, consisting of an enormous number of components, connected, interacting, and evolving via networks spanning multiple space and time scales. In traditional treatments of nervous-system networks, mechanoreceptor activity specifying the states of joints, muscles, and tendons flow through the spinal neurons to the brain. The challenge for this treatment is how the central executive can organize spatiotemporally distinct afferent signals to infer states of the whole body, segments, and appendages and to actively generate appropriate efferent signals. This challenge—called the “degrees of freedom” problem—is only compounded by the ambiguity and context-sensitivity of motor-unit and mechanoreceptor activity. This problem follows from a crucial premise about how this network divides its labor, that is, with strictly local processing at the periphery and global processing reserved for the center.

However, besides and cooperating with the central nervous system (CNS), other bodywide networks supporting perception allow peripheral and central processes to have an equal share in global coordination. Underneath our skin, a vast network of connective tissues and extracellular matrix (ECM) has been imagined as a multifractal tensegrity (MFT) in which the components hang together under tensional and compressional forces at multiple scales. This balance of tensional and compressional forces might offset local mechanical disturbances through the global realignment of forces [6–11], producing perceptual information ranging from coarse to fine [12–14]. If effortful perception is founded on action, then MFT-like cross-scale interactions proceeding through connective tissue may provide the biophysical substrate for perception of the body, attachments to the body, and surfaces adjacent to the body via dynamic touch [15,16]. Such networks support an “ultrafast” propagation of mechanical perturbations across vast distances called “preflex”, a faster-than-reflex response based on mechanical tensions rather than neural transmissions [17,18]. The situation of preflexes in the connective-tissue network’s self-similar, scale-free, fractal organization may resolve the degrees-of-freedom problem and support the spatiotemporal organization of afferent activity [15,16].

Testing whether bodywide MFT supports dynamic touch requires a specific analytical framework. Capacity for cross-scale interactions suggests the appearance of fractal organization that should support perceptual responses [15,19,20]. Indeed, fractal fluctuations of exploratory movements across the body [e.g., in hand, foot, head and postural center of pressure (henceforth, CoP)] all support the use of available mechanical information for generating perceptual judgments via dynamic touch [21–26]. The predictive role of fractal fluctuations appears to even extend across the body. When people manually heft a grasped object with their hands, the relatively distant measure of postural sway CoP at their feet has a fractal signature that helps predict the perceptual judgments [27,28]. Hence, fractal fluctuations provide a window into how specific patterns of movements spread across the entire body to support perceptual goals that seem—intuitively at least—specifically localized amidst the anatomical periphery.

### 1.3 Could flow of multifractal fluctuations support perception via dynamic touch?

Effortful manual wielding of an object prompts feedback loops spreading across the body at multiple spatiotemporal scales, rippling outwards from the wielding hand to the feet, maintaining an upright posture. These loops do not unfold in parallel at separate scales but rather interweave and intermix with each other, generating a nonlinear web of fluctuations throughout the body [23,25,26,29]. These nonlinearities generate multiple fractal forms following no less from spatial hierarchies of connective-tissue and neural networks than from the contextual constraints shaping action over time. Fractal fluctuations at any point in the body might spread through the rest of the body like contagion and this multifractal spread through the body matters for shaping perceptual judgments. Indeed, the bodywide flow of fractality indexes the flow of afferent information used to derive perceptual judgments for manually-hefted, visually-occluded objects, predicting individual differences in perceptual judgments from individual differences in bodywide flows of fractal fluctuations [30].

Hence, the human body is not a single point-mass that can be approximated by one fractal (i.e., monofractal) form. Instead, the body’s many degrees of freedom can each take on different monofractal forms. Our previous work used causal network modeling via vector autoregressive (VAR) analysis [31] to model pairwise exchanges of fractal fluctuations across 13 anatomical locations on the body and a handheld object. This approach opened a novel view of the human body as a multifractal field in which each degree of freedom might carry its single fractal form and through which individual degrees of freedom can influence others’ monofractal form. So, this portrait of the body is only multifractal in the sense that there are multiple monofractal forms spread across the body, opening up the capacity for fractal fluctuations to flow and change across the body.

Our previous work of developing a causal network of monofractal fluctuations was the first step. We now realize that this view was limited: rather than casting the body as a point-mass of one fractal form, it took a view of the body as a set of point masses, one for each degree of freedom. This higher-resolution view revealed preliminary insights, but it left the view of the body still relatively granular and insufficiently fluid; it was multifractal only at the macroscale of the whole body. If fractal fluctuations “flow” within one degree of freedom flexibly forcing or absorbing fluctuations on/from another, then the previous examination of how monofractal forms change across the body could only have been an initial step. Degrees of freedom are not point masses and may themselves contain finer-grained fluctuations supporting the bodywide flow.

Here, we aim to revisit this notion of a multifractal bodywide network with the recognition that a single component can itself be multifractal. That is, a single degree of freedom can exhibit different fractal patterns across time or for different-sized events. For that matter, perceptual accuracy may depend sooner on the nonlinearity generating multifractal forms than on nonlinearity generating monofractal forms [24,32–34]. The difference here has to do with the fact that monofractal form is only suggestive of a similar pattern observable at many scales, and it is mute to the reasons for similarity. Meanwhile, the multifractal form is explicitly the result of nonlinearities that force interactions across scales and not just coincidental resemblance of parallel but separate mechanisms [35].

The tensegrity proposal rests explicitly on the implication of interactions across scales, and thus the present multifractal revision of our previously monofractal results is an attempt to bring the evidence closer into alignment with the theory. The feedback loops in bodywide nonlinearities noted above reflect as well that feedback loops carry information amongst local and global scales: for example, local feedback loops unfolding amongst muscles of the hand both feed on and support more global feedback loops built between hand and legs planted on the ground. Such local-to-global and global-to-local flow of nonlinearity entails that each degree of freedom will be shimmering with multifractal form. Our previous work depicted the body as multifractal in the weak sense of spatially heterogeneous monofractal forms across the body. It construed the multifractal tensegrity as very many monofractal point masses bumping up against each other. However, if interactions across scales support perception, then it is important to elaborate prior work from monofractal analyses to multifractal analysis. Doing so will allow a more direct test of two points: 1) whether exchanges amongst individual degrees of freedom deals in fully multifractal fluctuations (i.e., not just spatially different monofractal fluctuations) and 2) whether the exchanges of multifractal fluctuations amongst degrees of freedom support the accuracy of perceptual judgments.

## 2. Materials and methods

### 2.1. Participants

Fifteen healthy adults (seven women, *mean*±*s.d.* age = 23.4±3.4 years, all right-handed [36]) with no muscular, orthopedic, and neurological disorder participated in this study after providing verbal and written informed consent.

### 2.2. Experimental objects

Each object (*n* = 6) consisted of an oak, hollow aluminum, or solid aluminum dowel (*l*×*d* = 1.2×75.0 cm; *m* = 68 g, 109 g, and 266 g, respectively) weighted by 4 or 12 stacked steel rings (*h* = 0.8 or 2.4 cm; *m* = 56 or 168 g; *d*_inner_ = 1.4 cm, *d*_outer_ = 3.4 cm) attached at 20.0 or 60.0 cm, respectively (table 1, figure 1*a*). The objects systematically differed in their mass, *m* (Object 1 > Object 2, Object 3 > Object 4, Object 5 > Object 6), the static moment, **M** (Object 1 = Object 2 = **M**_S_ < Object 3 = Object 4 = **M**_M_ < Object 5 = Object 6 = **M**_L_), and the moment of inertia, *I*_1_ and *I*_3_, reflecting the resistance of the object to rotation about the longitudinal axis (*I*_1_ values: Object 1, Object 2, Object 3 < Object 4, Object 5 < Object 6). A cotton tape of negligible mass was enfolded on each dowel to prevent the cutaneous perception of its composition.

**Table 1.**
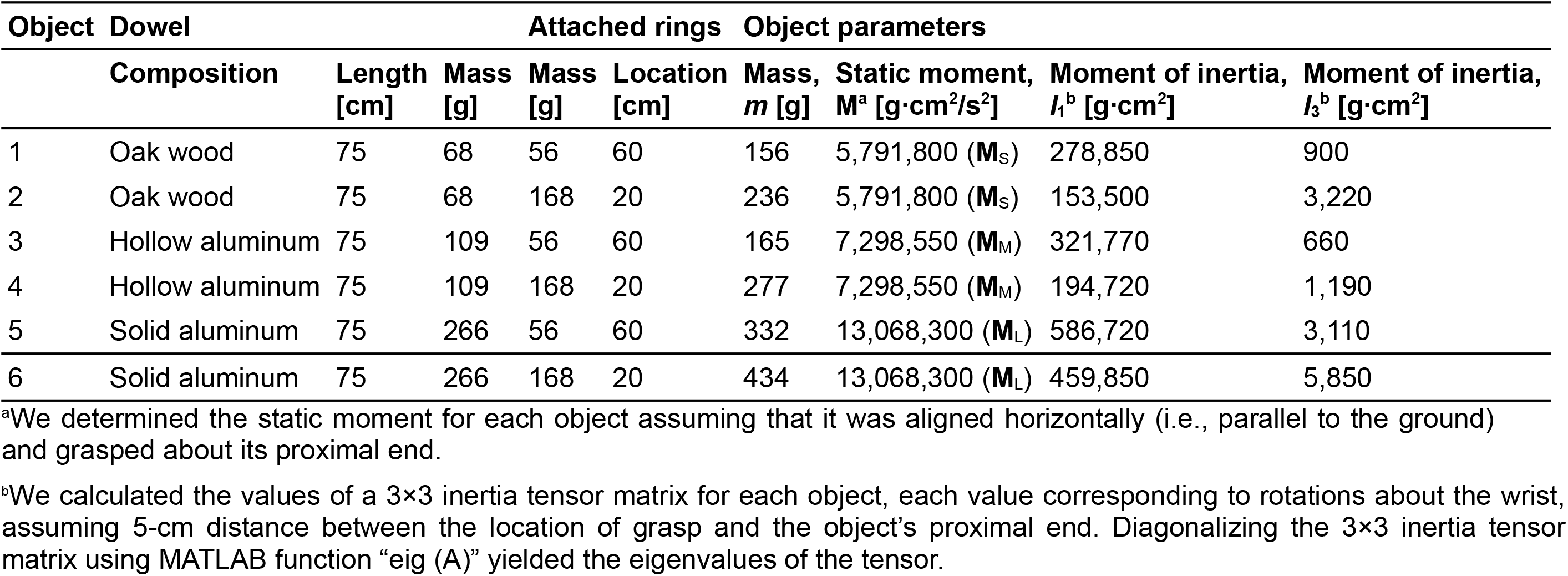
Experimental objects.

**Figure 1.**
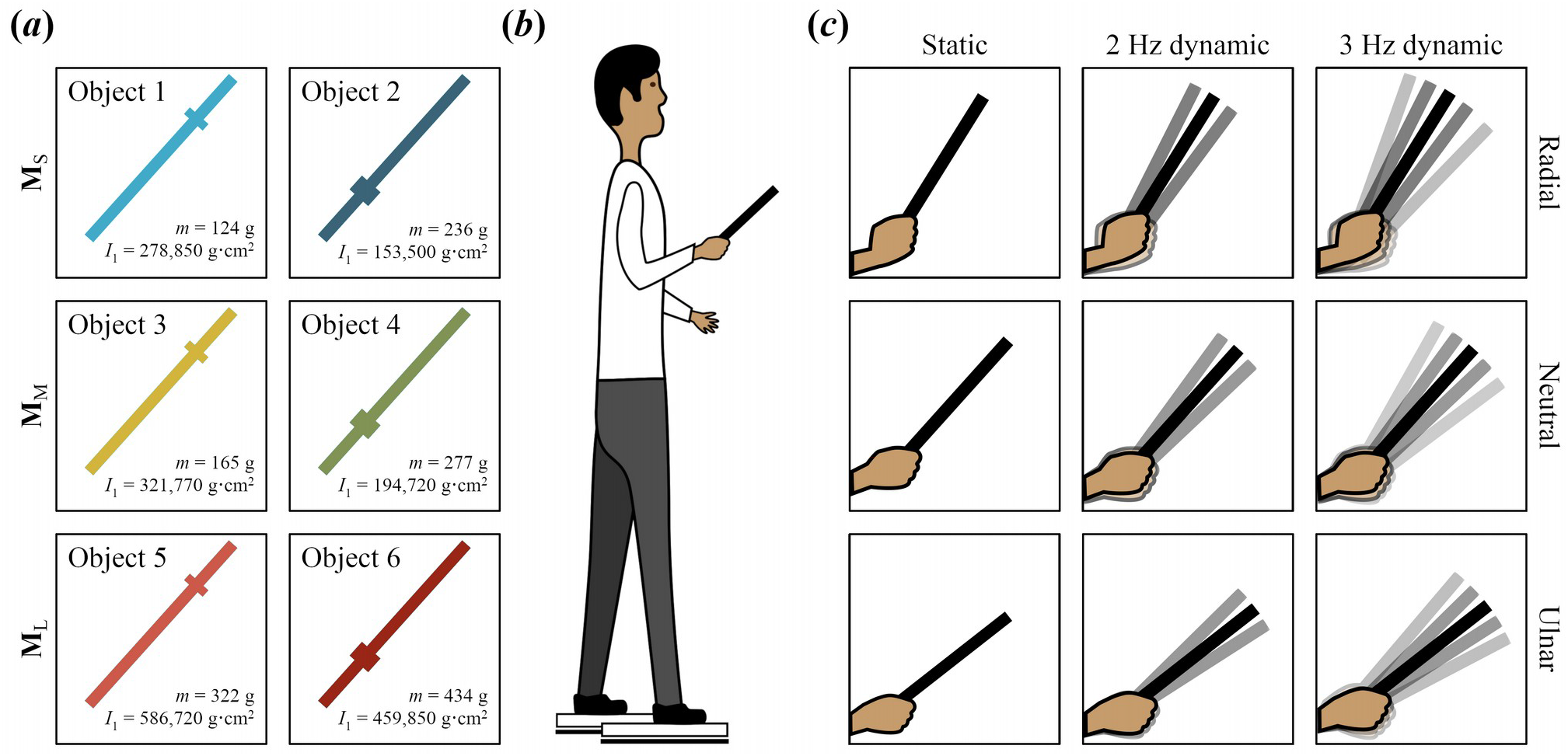
Schematic illustration of the experimental objects, setup, and exploratory conditions. (*a*) Each participant wielded six objects with different mass, *m*, static moment, **M**, and the moment of inertia, *I*_1_. (*b*) Each participant stood with his/her two feet on separate force plates, wielded each object for 5 s, and reported his/her judgments of heaviness and length of that object. (*c*) Different conditions of wrist angle and wrist angular kinematics.

### 2.3. Experimental setup and procedure

Each blindfolded participant stood on a pair of force plates (60×40 cm; Bertec Inc., Columbus, OH), wielded each object and reported judgments of heaviness and length (figure 1*b*). To impose constraints on haptic exploration, each participant moved his/her wrist about 10° ulnar deviation, the neutral position, or 10° radial deviation (figure 1*c*). In a static condition, the participant lifted and held each object static. In two dynamic conditions, the participant lifted and wielded each object synchronously with metronome beats at 2 Hz or 3 Hz, which added additional constraints on perceptual exploration. Each participant was instructed to minimize torso and upper-hand motion and the amplitude of wielding.

3D motion of three reflective markers (*d* = 9.5 mm) attached on each object at 30, 45, and 60 cm and nine reflective markers attached on the participant’s body (table 2, figure 2*a*) was tracked at 100 Hz using an eight-camera Qualisys motion-tracking system (Qualisys Inc., Boston, MA).

**Table 2.**
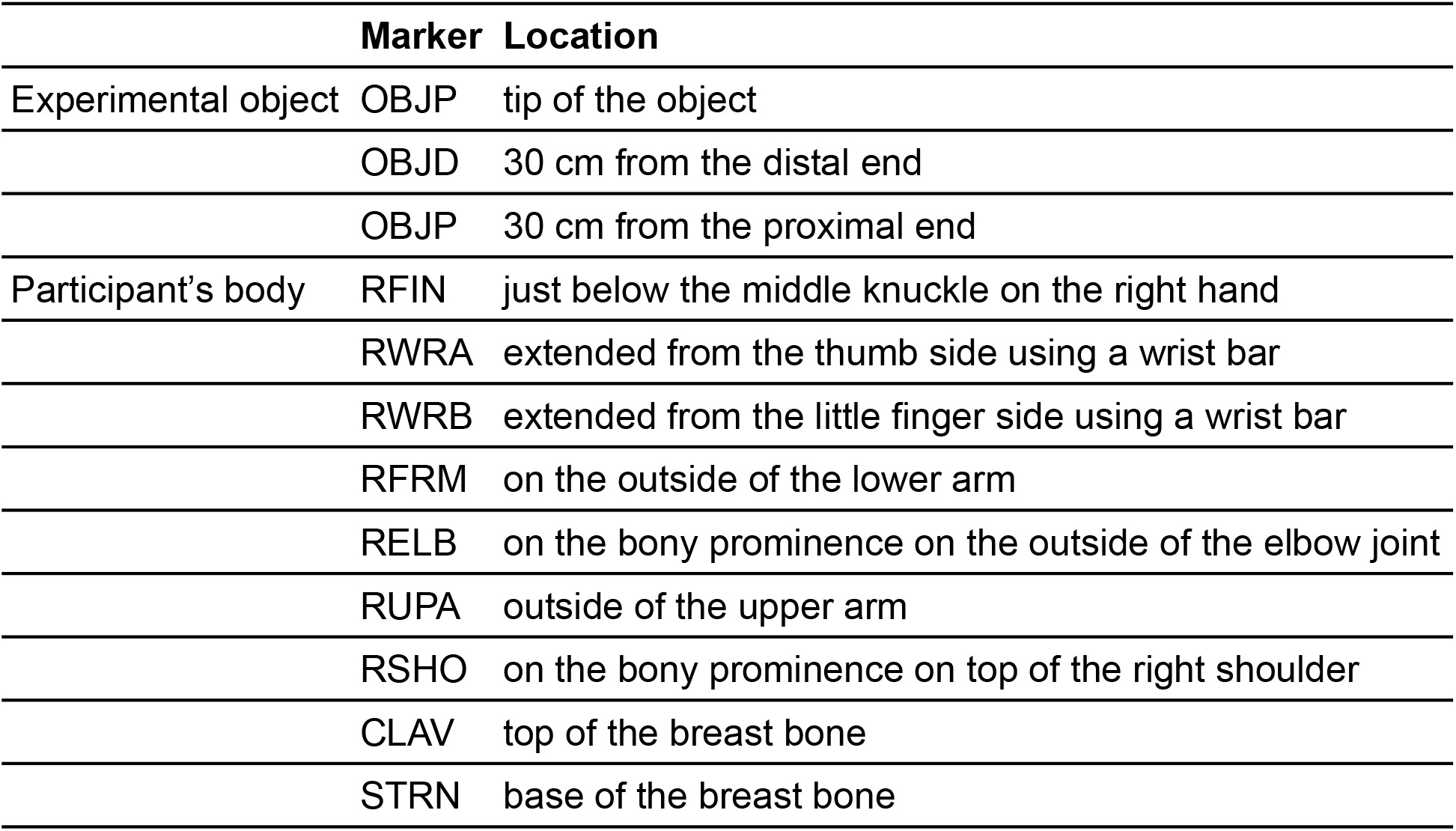
Location of the reflective markers attached to each experimental object and the participant’s body.

**Figure 2.**
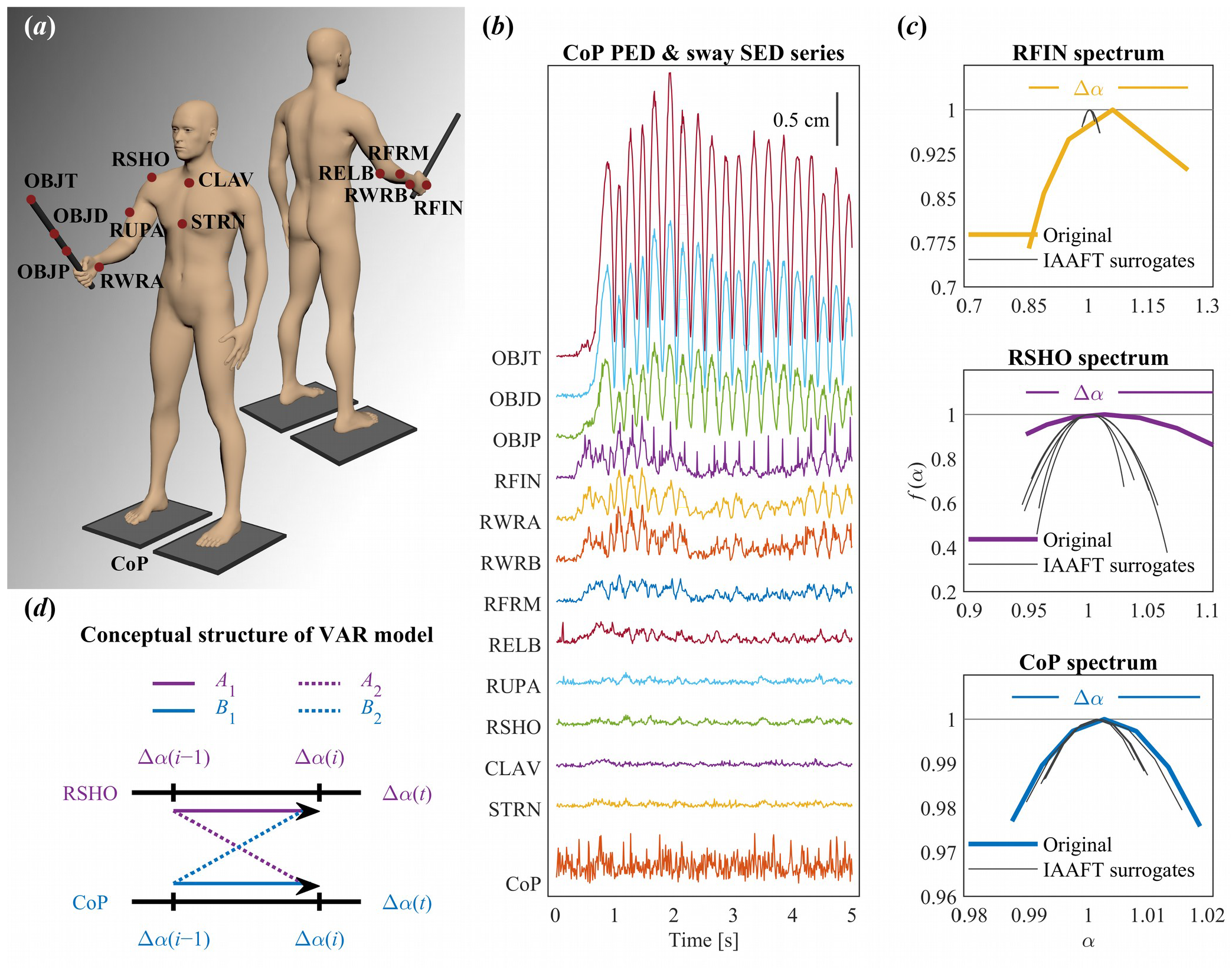
Overview of data acquisition process and analysis. (*a*) Locations of the reflective markers attached to the experimental object and the participant’s body. (*b*) CoP PED and sway SED series for a representative trial (Condition: Neutral, 2 Hz dynamic, Object 6). (*c*) Singularity spectrums (*α*, *f*(*α*)) of a representative original CoP PED series and two sway SED series (colored lines), as well as those of their five IAAFT surrogates (gray lines). (*d*) The conceptual structure of the VAR analysis used to model the diffusion of multifractality across different anatomical locations. The contribution of each location is represented as a series of trial-by-trial values of the singularity spectrum width (*Δ α* =*α_max_* −*α_min_*). Arrows represent weights in the model, indicating the effects of *Δ α* in the previous trail on *Δ α* in the current trial.

Each participant completed 108 trials (3 Wrist angles × 3 Wrist angular kinematics × 6 Objects × 2 Trials/Object). Each factor of Wrist angle (Radial, Neutral and Ulnar) was crossed with each factor of Wrist angular kinematics (Static, 2 Hz dynamic and 3 Hz dynamic). The order of the 12 trials (6 Objects × 2 Trials/Object) was pseudo-randomized for each block. Before the first and after every six trials, each participant wielded a reference object that was arbitrarily attributed to a heaviness value of 100 units. In each trial, after a “lift” signal, the participant lifted the object by about 5 cm and held it static or wielded it at 2 Hz or 3 Hz. After 5 s, following a “stop” signal, the participant placed the object back and reported (a) perceived heaviness proportionally higher and lower than 100 to an object perceived heavier and lighter, respectively, than the reference object; and (b) perceived length by adjusting the position of a marker on a string-pulley assembly.

### 2.4. Data processing

#### 2.4.1. CoP planar Euclidean displacement (PED) series

Force plate output was downsampled by 1/20 (i.e., from 2000 Hz to 100 Hz) to match motion-tracking sampling rates. The ground reaction forces recorded at each trial yielded a two-dimensional CoP series of 500 samples, describing the position of the CoP along the participant’s medial-lateral and anterior-posterior axes. A one-dimensional CoP planar Euclidean displacement (PED) series of 499 samples was obtained for each downsampled CoP series, describing CoP displacement along the transverse plane of the body (figure 2*b*).

#### 2.4.2. Sway spatial Euclidean displacement (SED) series

Motion tracking of each reflective marker (*n* = 12) yielded a three-dimensional kinematic series of 500 samples, describing its position along the participant’s medial-lateral, anterior-posterior and superior-inferior axes. A one-dimensional spatial Euclidean displacement (SED) series of 499 samples was obtained for each marker describing the displacement of that marker in 3D (figure 2*b*).

### 2.5. Assessing multifractality and interactivity

#### 2.5.1. Direct estimation of multifractal spectrums

Chhabra and Jensen’s direct method estimated multifractal spectrums of CoP PED and sway SED series [37]. This method samples series *u* (*t*) at progressively larger scales such that the proportion of signal *P_i_* (*L*) falling within the *i^th^* bin of scale *L* is

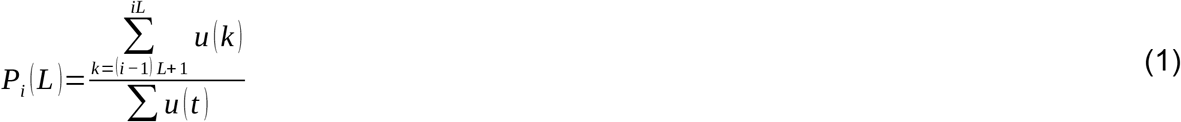

As *L* increases, *P_i_* (*L*) represents progressively larger proportion of *u* (*t*),

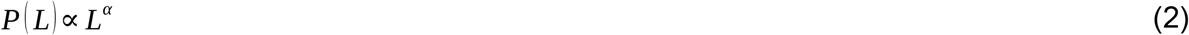

suggesting growth of proportion according to one “singularity” strength *α* [38]. *P* (*L*) exhibits multifractal dynamics when it grows heterogeneously across time scales *L* according to multiple singularity strengths, such that

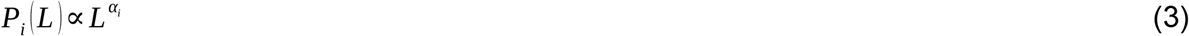

whereby each *i^th^* bin may show a distinct relationship of *P* (*L*) with *L*. The width of this singularity spectrum, *Δ α*(*α_max_* − *α_min_*), indicates the heterogeneity of these relationships [39,40].

Chhabra and Jensen’s method [37] estimates *P* (*L*) for *N_L_* nonoverlapping bins of *L*-sizes and transforms them into a “mass” *μ* using a *q* parameter emphasizing higher or lower *P* (*L*) for *q* >1 and *q* <1, respectively, as follows

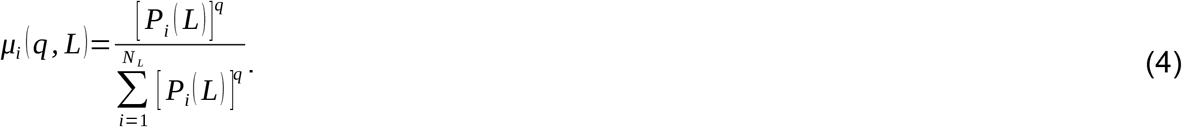

*α* (*q*) is the singularity for mass μ(q)-weighted P(L) estimated by

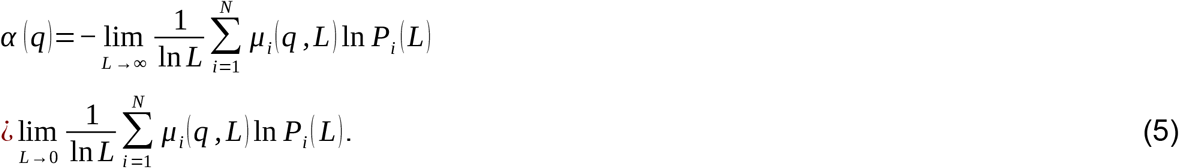

Each estimated value of *α* (*q*) belongs to the singularity spectrum only when the Shannon entropy of *μ* (*q, l*) scales with *L* according to the Hausdorff dimension *f* (*q*), where

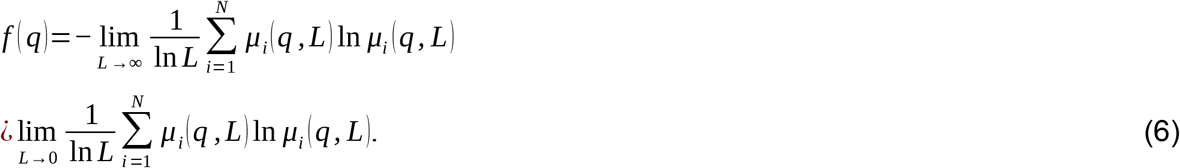

For values of *q* yielding a strong relationship between Eqs. (5 & 6)—in this study, exhibited a correlation coefficient, *r* > 0.9, the parametric curve (*α* (*q*) *, f* (*q*)) or (*α, f* (*α*)) constitutes the singularity spectrum. The singularity spectrum width, *Δ α* =*α_max_* −*α_min_*, was calculated for each CoP PED and sway SED series (figure 2*c*).

#### 2.5.2. Surrogate testing

To identify whether nonzero *Δ α* reflected nonlinear temporal correlations [41,42], *Δ α* of each original series was compared to *Δ α* of surrogate series using Iterated Amplitude Adjusted Fourier Transformation (IAAFT) [43,44]. IAAFT randomizes original values time-symmetrically around the autoregressive structure. It thus generates surrogates that randomize phase ordering of the series’ spectral amplitudes while preserving only linear temporal correlations. If the original *Δ α* exceeds a 95% confidence interval (CI) of *Δ α* for 32 IAAFT series (i.e., *p* < 0.05), then the original time series has nonzero nonlinear correlations quantifiable using the one-sample t-statistic (henceforth, *t*_MF_) comparing *Δ α* for the original series to that for the surrogates.

### 2.6. Vector autoregression (VAR) analysis

Vector autoregression (VAR) captures linear interdependencies amongst concurrent series. We used VAR to model the effects of trial-by-trial *Δ α* from each marker position to each other marker position across trials in the study. VAR describes each variable based on its own lagged value and that of each other variable, along with an error term. Unlike structural models, VAR modeling does not use any prior knowledge besides a list of variables that can be hypothesized to affect each other temporally. Thus, VAR analysis allows for exploring causal relationships after addressing minimal short-lag relationships [45].

VAR produces a system of *m* regression equations predicting each variable as a function of lagged values of itself and of each other variable. In the simplest case of *m*=*2*, with time series *f* (*t*) and *g* (*t*) definable at each value of time *t* =1 to *t* =*N*, the structure of a VAR model is:

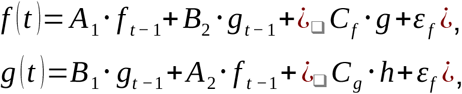

where *A_j_* and *B_j_* quantify the effects of the previous values of *f* and *g*, respectively, with *j* indexing the variable to which these previous values contribute, with error terms *ε_f_* and *ε_g_* [46]. The above equations describe a 1-lag VAR. Each *f* and *g* is described in terms of values up to 1 value preceding the predicted values. VAR models can include exogenous variables, such as the factors of experimental design which stand outside the relationships amongst the variables internal to the system. In the above example, the time series *h* (*t*) can induce changes in *f* (*t*) or *g* (*t*), but changes in neither *f* (*t*) or *g* (*t*) can induce changes in *h* (*t*). *h* is an exogenous variable, and *C_f_* and *C_g_* are quantify the effects of *h* (*t*) on *f* (*t*) and *g* (*t*), respectively. Endogenous variables are variables internal to the system (i.e., *f* (*t*) or *g* (*t*)) which may respond to and induce changes in other endogenous variables. For the present analysis, *Δ α* of CoP PED and sway SED series served as an endogenous variable (figure 2*d*).

VAR models forecast the effects of endogenous variables through impulse-response functions (IRFs). As opposed to standard regression, which can only evaluate the relationship between *f* (*t*) and *g* (*t*), IRFs can evaluate the relationship between *f* (*t*) and *g* (*t* + *τ*), or between *g* (*t*) and *f* (*t* +*τ*), where *τ* is a whole number. First, orthogonalizing the regression equations and, second, inducing an “impulse” to the system of regression equations by adding 1 *s.e.m.* to any single variable, propagating responses across variables. The plot of an IRF describes how an impulse in one time series changes the later predicted values in a different time series [46,47]. It is customary to fit the least number of lags that leave independently and identically distributed residuals.

### 2.7. Statistical analysis

All pairwise impulse-response relationships indicated the effects of increases in *Δ α* across subsequent trials. A full-factorial regression model [48] of Impulse × Response × Trial tested the average effects and responses of each marker position along with orthogonal linear, quadratic, and cubic polynomials of Trial, using the “nlme” package for RStudio [49]. Impulse and Response served as class variables indicating the locations of the impulse variables and the responding variables, respectively.

Regression models of unsigned error, for example, absolute(*H*/*L*_perceived_ − *H*/*L*_actual_), tested the effects of significant impulse-response relationships between marker positions. Unsigned error for *L*_perceived_ was the absolute value of *L*_perceived_ minus actual length (i.e., 75 cm). Because *H*_perceived_ was calculated as a percentage rating relative to a 109-g reference object, *H*_error_ was calculated as the absolute value of [*H*_actual_×(*H*_perceived_×109)/100−100] rounded to the nearest whole-number percentage. For instance, perceiving the heaviness of 236-g Object 2 as 120% entails unsigned error *H*_error_ = absolute(100×[236×(120×109)/100)]/236–100) = 45. Because *L*_error_ was linear (i.e., additive) and *H*_error_ was nonlinear (i.e., a rate), they were modeled using linear mixed-effect (LME) and mixed-effect Poisson regression, using “nlme” and “lme4” packages for Rstudio, respectively [49,50].

Predictors of unsigned error included Wrist angle, Object’s logarithmic moments of inertia (Log*I*_1_ and Log*I*_3_), trial order, *Δ α* of CoP (CoP_Δ*α*_), and the IRF values forecasting the first significant response to impulse over subsequent trials for OBJD->RFIN, OBJD->RELB, OBJD->RUPA, RWRA->RELB, RFIN->RUPA, RELB->RWRA, RELB->RUPA, RELB->RSHO, RUPA->CLAV, CoP->RWRA and RFRM->CoP. Log*I*_3_ and trial order improved model fit only for *H*_error_ and *L*_error_, respectively.

## 3. Results

### 3.1. Fluctuations at each anatomical location exhibits multifractality

All original CoP PED and sway SED series exhibited non-zero singularity-spectrum widths (i.e., *Δ α* > 0; range of *Δ α*: CoP: 0.0085–0.48; OBJT: 0.026–0.85; OBJD: 0.029–0.80; OBJP: 0.017–0.64; RFIN: 0.028–0.90; RWRA: 0.036–0.85; RWRB: 0.023–0.85; RFRM: 0.015–1.16; RELB: 0.028–1.41; RUPA: 0.033–1.81; RSHO: 0.024–2.09; CLAV: 0.033–1.89; STRN: 0.033–1.80). *Δ α* was wider for 1412 of 1620 CoP PED series, as well as for each original sway SED series than the corresponding IAAFT surrogates (figure 3).

**Figure 3.**
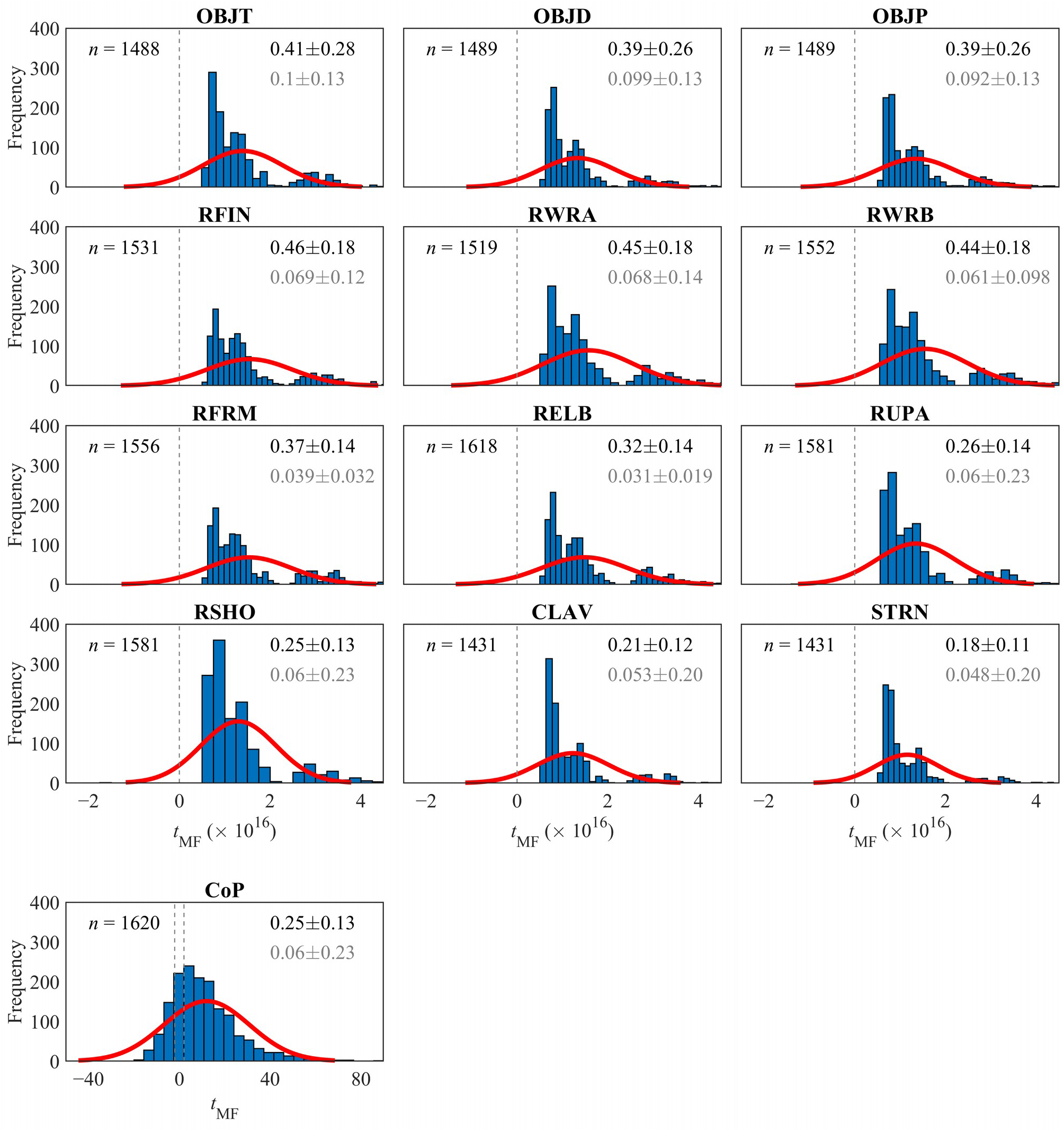
Frequency distributions of *t*_MF_ comparing the singularity spectrum widths (*Δ α* =*α_max_* −*α_min_*) of the original CoP PED and sway SED series and that of their 32 IAAFT surrogates. The values on the top right in black and gray in each plot describe *mean*±*s.d.* values of *Δ α* for the original version and 32 IAAFT surrogates of the recorded CoP PED and sway SED series the number of which is indicated on the top left. *t*_TM_ > 0 indicates that the original spectrum was wider than the surrogate spectrums and vice versa. The dashed vertical lines indicate the cutoffs for statistical significance at the two-tailed alpha level of 0.05 for 31 DoFs. Most (1412/1620) CoP PED and all sway SED series showed multifractality.

### 3.2. Multifractal fluctuations flow across the body

First, the IRFs showed pairwise exchanges of multifractality following the sequence of motor segments from a handheld object (OBJD) to the shoulder (RSHO; figure 4). The strongest effects included multifractality-promoting effects from the object (OBJD) on the most distal arm segments that become progressively smaller (from RFIN to RWRA to RFRM) and then multifractality-diminishing effects on the proximal arm segments (RELB, RUPA and RSHO). Hence, the local contacts with objects at hand have intuitive effects on the chain of motor degrees of freedom, with multifractal fluctuations decaying as they propagate from peripheral to central components. [The regression modeling confirmed that the individual mean differences from zero are significant (Supplementary Table S1).]

**Figure 4.**
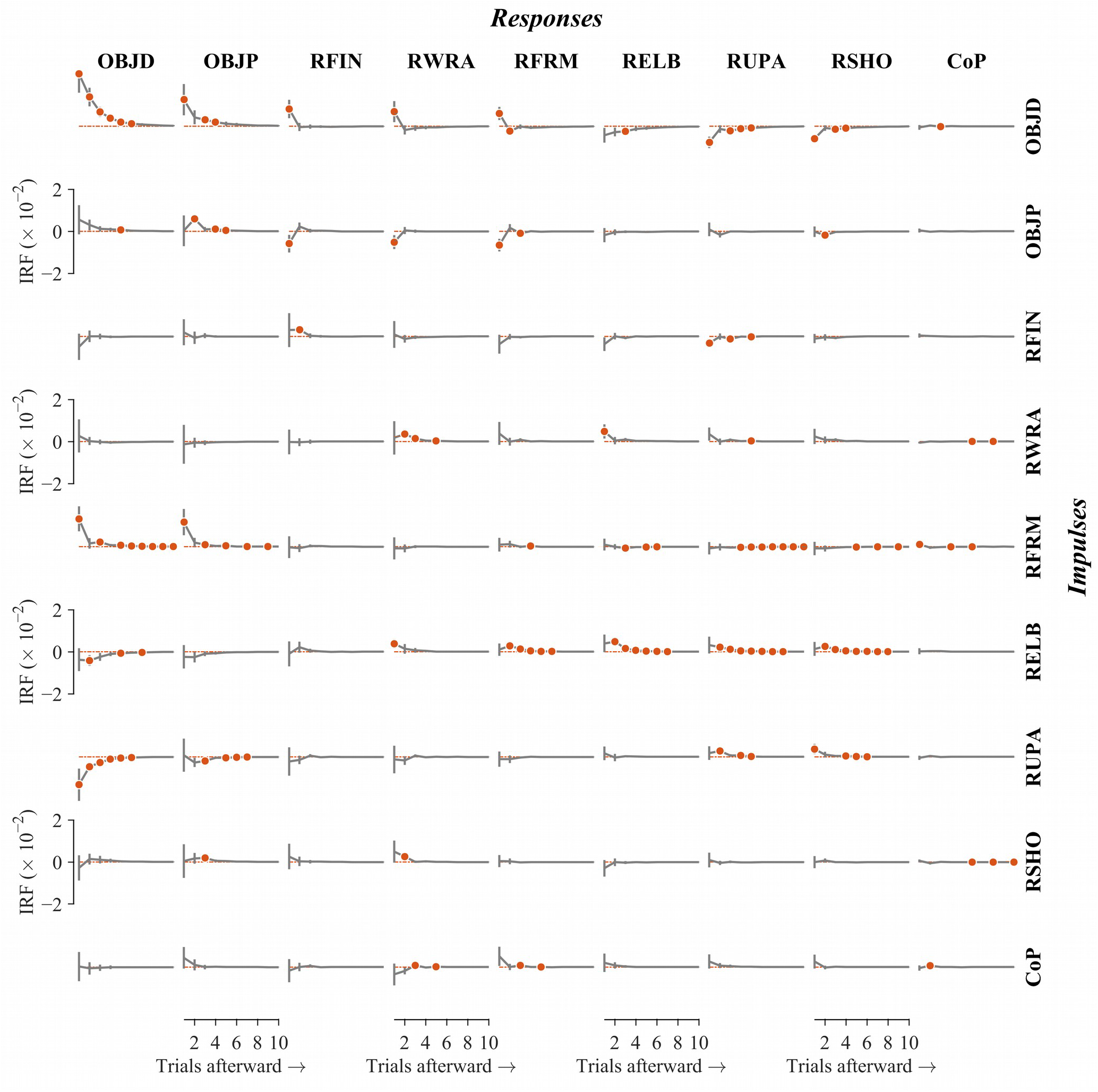
*Mean*±*s.e.m.* (*n* = 15 participants) responses in *Δ α* of CoP PED and sway SED series over ten trials afterward to an impulse in *Δ α* of each other series in the current trial. Each black curve illustrates the later response as it decays over subsequent trials, and each solid red circle indicates a significant (*p* < 0.01) response to an impulse in *i^th^* trial afterward (1 through 10). The strongest effects included multifractality-promoting effects from the object (OBJD) on the most distal arm segments that become progressively smaller (from RFIN to RWRA to RFRM) and then multifractality-diminishing effects on the proximal arm segments (RELB, RUPA and RSHO); see text for all other significant impulse-response effects.

Collectively, the IRF results suggest functional segregation of forearm from upper arm in how each mediated exchanges of multifractal fluctuations. The forearm sent multifractal fluctuations outward to the object, indicating that forearm and object promote each other’s multifractality. This mutual object-forearm promotion of multifractality came at the expense of upper-arm multifractality. Perhaps the forearm draws multifractality away from the upper arm to send downstream to the object. Certainly, increases in RFRM and RFIN multifractality decreased later RELB and RUPA multifractality, respectively. Conversely, increases in RUPA and RELB multifractality decreased later OBJD multifractality, with increases in RUPA also contributing to decreases in multifractality at OBJD.

The joints played intermediating roles between forearm-like multifractality-promoting and upper-arm-like multifractality-limiting tendencies. Exemplifying the former, OBJD and RELB multifractality decreased when the other increased, and increases in RELB multifractality prompted increases in multifractality across the upper arm (RUPA and RSHO). Exemplifying the latter, RELB and RWRA showed mutual positive effects on each other’s multifractality as though RELB might participate in the forearm’s support of multifractality. Similar to RELB, RSHO showed the upper arm’s inverse relationship to increases in OBJD multifractality and increased with RUPA multifractality in response to RELB multifractality. But RSHO also showed a multifractality-promoting aspect: increases in RSHO multifractality prompted later increases in OBJD and RWRA multifractality.

The IRF effects extended beyond the upper body to include CoP. Increases in CoP multifractality showed a positive effect on later RWRA and RFRM multifractality.

### 3.3. The flow of multifractal fluctuations across the body influences perceptual accuracy

The flow of multifractal fluctuations differed across individuals and predicted individual differences in accuracy. H_error_ depended on seven pairwise exchanges of multifractal fluctuations in supporting perceptual accuracy (table 3). The GLM returned positive coefficients for IR effects of OBJD on RFIN (*b*±*s.e.m.* = 10.54±1.99, *p* < 0.001), OBJD on RUPA (*b*±*s.e.m.* = 12.56±3.68, *p* < 0.001), RFIN on RUPA (*b*±*s.e.m.* = 20.75±6.42, *p* = 0.001), RELB on RUPA (*b*±*s.e.m.* = 15.36±5.62, *p* = 0.006) and RUPA on CLAV (*b*±*s.e.m.* = 10.10±4.61, *p* = 0.028). It returned negative coefficients for IR effects of OBJD on RELB (*b*±*s.e.m.* = –10.07±3.65, *p* = 0.006) and CoP on RWRA (*b*±*s.e.m.* = –51.54±11.51, *p* < 0.001). The negative IR effects of OBJD on RELB and CoP on RWRA entailed that these pairwise exchanges of multifractal fluctuations entailed decreases in *H*_error_; all other IR effects entailed increases in *H*_error_ (figure 5, left panels). These effects held above and beyond the significant effects of wrist angle variations, and of the logarithmic first and third moments of inertia (table 3). Also, CoP_Δ*α*_ showed a positive effect on *H*_error_ (*b*±*s.e.m.* = 0.59±0.14, *p* < 0.001), suggesting that greater *Δ α* led to less accurate heaviness judgments, as shown previously [28].

**Figure 5.**
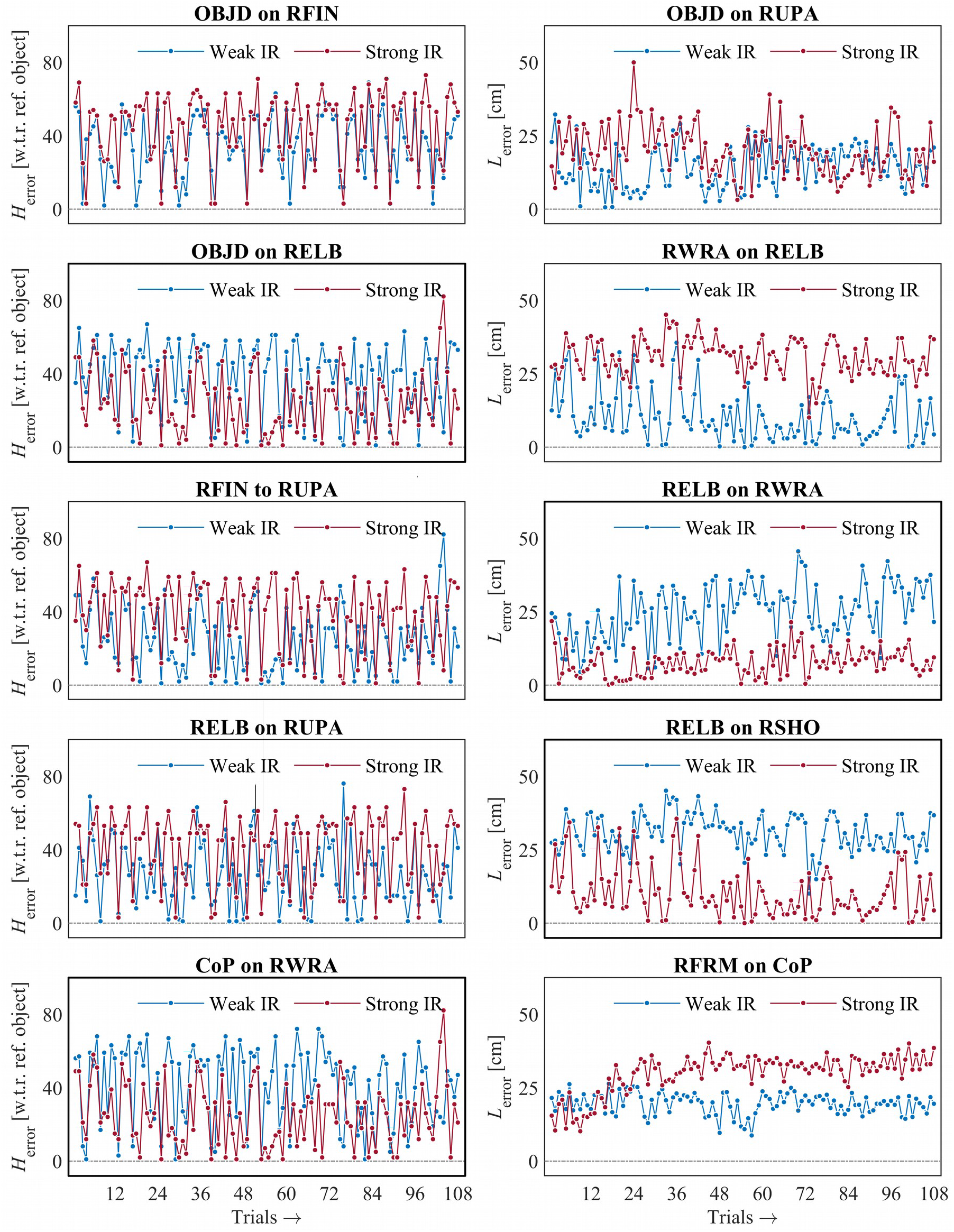
Comparisons of absolute errors in perceived heaviness, *H*_error_, and perceived length, *L_error_*, for representative participants with weak and strong impulse-response (IR) effects for selected pairwise relationships. The strong IR effects of OBJD on RELB and CoP on RWRA entailed decrease in *H*_error_ (left panels in bold); all other IR effects entailed increases in *H*_error_ (left panels). The strong IR effects of RELB on RWRA and RELB on RSHO entailed decrease in *L*_error_ (right panels in bold); all other IR effects entailed increases in *L*_error_ (right panels).

**Table 3.**
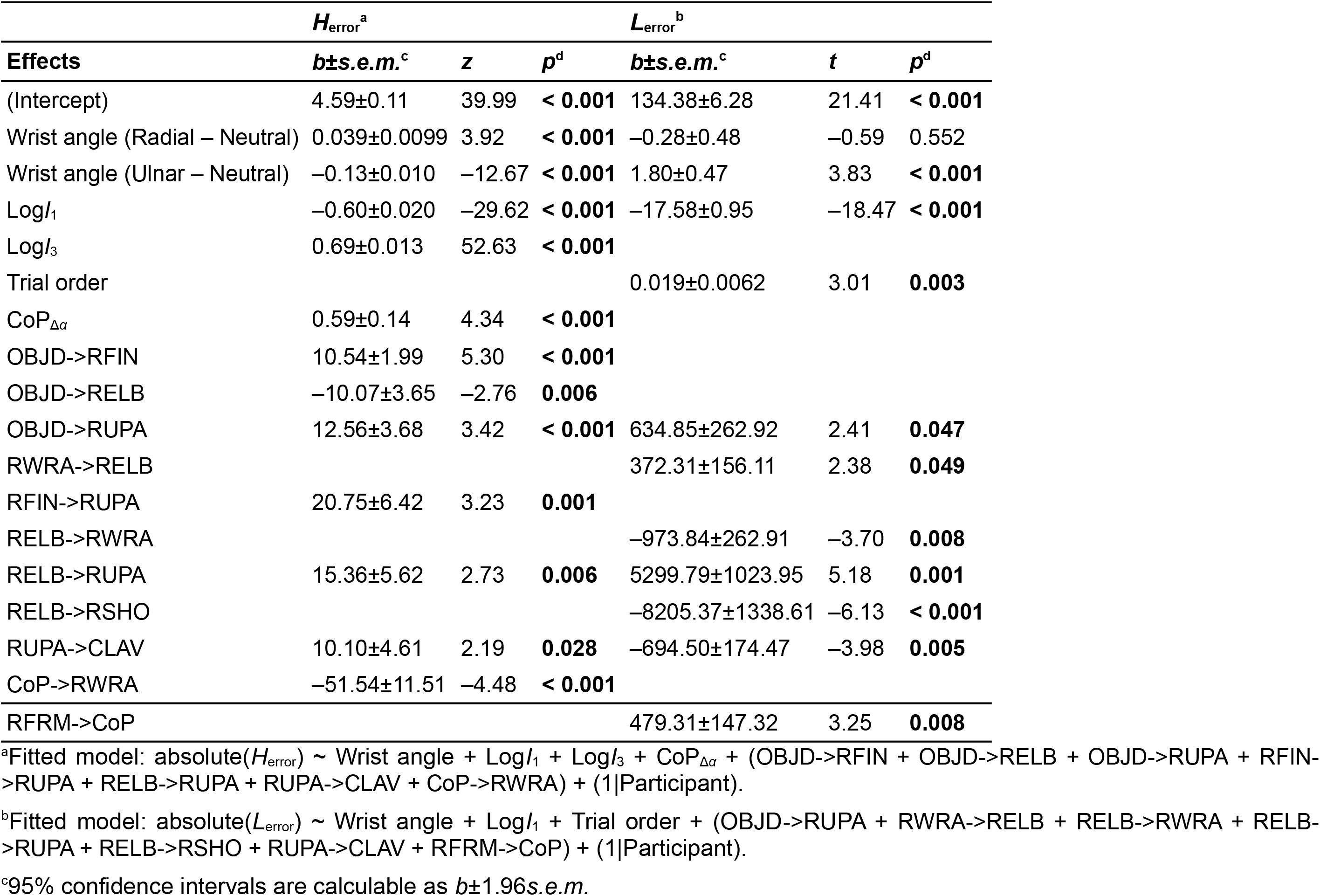
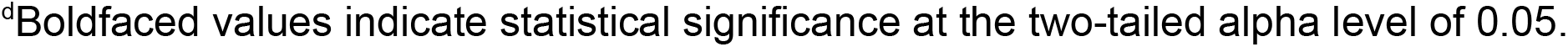
Coefficients of GLM and LME models examining the effects of CoP multifractality and significant impulse-response relationships on the absolute error in perceived heaviness, *H*_error_, and perceived length, *L*_error_, respectively.

| Effects | $H_{\text{error}}^a$ | | | $L_{\text{error}}^b$ | | |
| --- | --- | --- | --- | --- | --- | --- |
| | $b \pm \text{s.e.m.}^c$ | $z$ | $p^d$ | $b \pm \text{s.e.m.}^c$ | $t$ | $p^d$ |
| (Intercept) | 4.59±0.11 | 39.99 | < **0.001** | 134.38±6.28 | 21.41 | < **0.001** |
| Wrist angle (Radial – Neutral) | 0.039±0.0099 | 3.92 | < **0.001** | –0.28±0.48 | –0.59 | 0.552 |
| Wrist angle (Ulnar – Neutral) | –0.13±0.010 | –12.67 | < **0.001** | 1.80±0.47 | 3.83 | < **0.001** |
| Log$I_1$ | –0.60±0.020 | –29.62 | < **0.001** | –17.58±0.95 | –18.47 | < **0.001** |
| Log$I_3$ | 0.69±0.013 | 52.63 | < **0.001** | | | |
| Trial order | | | | 0.019±0.0062 | 3.01 | **0.003** |
| CoP$_{\Delta\alpha}$ | 0.59±0.14 | 4.34 | < **0.001** | | | |
| OBJD->RFIN | 10.54±1.99 | 5.30 | < **0.001** | | | |
| OBJD->RELB | –10.07±3.65 | –2.76 | **0.006** | | | |
| OBJD->RUPA | 12.56±3.68 | 3.42 | < **0.001** | 634.85±262.92 | 2.41 | **0.047** |
| RWRA->RELB | | | | 372.31±156.11 | 2.38 | **0.049** |
| RFIN->RUPA | 20.75±6.42 | 3.23 | **0.001** | | | |
| RELB->RWRA | | | | –973.84±262.91 | –3.70 | **0.008** |
| RELB->RUPA | 15.36±5.62 | 2.73 | **0.006** | 5299.79±1023.95 | 5.18 | **0.001** |
| RELB->RSHO | | | | –8205.37±1338.61 | –6.13 | < **0.001** |
| RUPA->CLAV | 10.10±4.61 | 2.19 | **0.028** | –694.50±174.47 | –3.98 | **0.005** |
| CoP->RWRA | –51.54±11.51 | –4.48 | < **0.001** | | | |
| RFRM->CoP | | | | 479.31±147.32 | 3.25 | **0.008** |

<sup>a</sup>Fitted model: $\text{absolute}(H_{\text{error}}) \sim \text{Wrist angle} + \text{Log}I_1 + \text{Log}I_3 + \text{CoP}_{\Delta\alpha} + (\text{OBJD->RFIN} + \text{OBJD->RELB} + \text{OBJD->RUPA} + \text{RFIN-}$ $\text{>RUPA} + \text{RELB->RUPA} + \text{RUPA->CLAV} + \text{CoP->RWRA}) + (1|\text{Participant})$ .
<sup>b</sup>Fitted model: $\text{absolute}(L_{\text{error}}) \sim \text{Wrist angle} + \text{Log}I_1 + \text{Trial order} + (\text{OBJD->RUPA} + \text{RWRA->RELB} + \text{RELB->RWRA} + \text{RELB-}$ $\text{>RUPA} + \text{RELB->RSHO} + \text{RUPA->CLAV} + \text{RFRM->CoP}) + (1|\text{Participant})$ .
<sup>c</sup>95% confidence intervals are calculable as $b \pm 1.96\text{s.e.m.}$
<sup>d</sup>Boldfaced values indicate statistical significance at the two-tailed alpha level of 0.05.

L_error_ depended on seven pairwise exchanges of multifractal fluctuations in supporting perceptual accuracy. The LME returned positive coefficients for IR effects of OBJD on RUPA (*b*±*s.e.m.* = 634.85±262.92, *p* = 0.047), RWRA on RELB (*b*±*s.e.m.* = 372.31±156.11, *p* = 0.049), RELB on RUPA (*b*±*s.e.m.* = 5399.79±1023.95, *p* = 0.001) and RFRM to CoP (*b*±*s.e.m.* = 479.31±147.32, *p* = 0.008). It returned negative coefficients for IR effects RELB on RWRA (*b*±*s.e.m.* = –973.84±262.91, *p* = 0.008), RELB on RSHO (*b*±*s.e.m.* = – 8205.37±1338.61, *p* < 0.001) and RUPA on CLAV (*b*±*s.e.m.* = –694.50±174.47, *p* = 0.005). Thus, *L*_error_ decreased with the transfer of multifractal fluctuations from RELB to RWRA, RELB to RSHO and RUPA to CLAV, and *L*_error_ increased with the transfer of multifractal fluctuations from OBJD to RUPA, RWRA to RELB, RELBV to RUPA and RFRM to CoP (figure 5, right panels). These effects held above and beyond significant effects of wrist angle variations and of the logarithmic first moment of inertia (table 3).

## 4. Discussion

We investigated whether and how the flow of multifractal fluctuations entailed in the bodywide MFT supports perception via dynamic touch. We expected that if perception via dynamic touch occasions an upstream flow of information from the point of distal stimulation (i.e., the hand), which sources multifractality from the global dynamics, then not only should multifractality at hand affect multifractality at the lower and upper arms (i.e., reflecting upstream effects of distal hand activity) but also multifractality in CoP should promote multifractality at hand (i.e., posture is at the forefront of multifractality resources for the distal body parts). Our findings support this hypothesis.

The observed multifractality was due to nonlinear interactions across scales reflecting feedback loops proceeding locally, globally, and interacting across the scales. The impulse-response forecasting obtained from VAR analysis revealed upstream effects of the distal hand activity, as multifractal fluctuations at hand promoted multifractal fluctuations at the lower arm segments and reduced it in the upper arm segments. Multifractality in the global measure of CoP helped promote multifractal fluctuations at hand. The strength of these exchanges of multifractal fluctuations amongst degrees of freedom indexed the accuracy of perception. These results strengthen the view that nonlinear interactions entailed by the bodywide MFT support the flow of mechanical information supporting the coordination of perceptual judgments of object heaviness and length [30].

Collectively, our results offer a window into the bodywide synergy supporting dynamic touch by the hand. Multifractality-promoting effects of OBJD on the most distal parts of the arm became progressively smaller (from RFIN to RWRA to RFRM), and multifractality-diminishing effects of OBJD extend along with the proximal parts of the arm towards the shoulder (RELB, RUPA and RSHO). The joints thus played a mediating role between the upper arm and forearm; for instance, RELB showed a multifractality-limiting effect on OBJD and a multifractality-promoting effect on RWRA, RUPA and RSHO. And although RSHO showed increases in multifractality in conjunction with that of the rest of the upper arm, RSHO broke ranks with the upper arm and promoted later RWRA multifractality. Finally, increases in CoP multifractality precede subsequent increases in both RWRA and RFRM multifractality, situating local fluctuations at hand into a global context.

Our models of absolute error showed that perceptual accuracy in dynamic touch hinges on specific flow of multifractal fluctuations across the body. The strength of IRF effects served as significant predictors of the absolute error in judgments of both heaviness and length. Greater flow of multifractality across all pairs of anatomical locations did not always resulted in more accurate judgments, but the flow of multifractal fluctuations across specific body segments played a crucial role in perceiving accurately. Hence, we cannot claim the simple wholesale conclusion that more multifractality entails higher accuracy [27,28]. Instead, dynamic touch hinges upon specific interplay amongst many degrees of freedom, each individually fluctuating multifractally—that is, with multiple fractal forms across time and fluctuation size—and the flow of these multifractal fluctuations may provide an essential medium for perceptual information [12,14].

Our findings strengthen the emerging view that a wider-than-neural set of tissues enable preflexes, that is, mechanotransduction of contextually-specific responses flowing faster than neural transmission. Crucially, the concept of “preflexes” appears to be more generic to bodywide coordination than specific to local anatomical structures [51–53]. Indeed, if the MFT is the architecture of life [54], then preflexes must be foundational to how life perceives and acts.

## 5. Conclusion

Our findings make a compelling case that the study of perception might not be exhausted by activity in the CNS. Instead, it must also include the flow of multifractal fluctuations across the bodywide MFT. Indeed, far from suggesting the latter to the exclusion of the former, it is incredibly likely that the CNS and MFT are mutually supporting systems [15]. The network relationships we have presented across the anatomical sleeves of the body show close resemblance to the resting state network (RSN) dynamics exhibited by the central nervous system (CNS) [55,56]. What RSN dynamics proposes for networks of neurons, we suggest the existence of synergies specific to perceptual intent (e.g., object heaviness vs. length vs. shape) in the flow of multifractality fluctuations in the network of anatomical nodes across the body.

Future research into the endogenous and exogenous factors affecting the bodywide flow of multifractal fluctuations might support diverse clinical applications. For instance, fractal fluctuations in exploratory movements predict differences in dynamic touch capabilities between children with typical and atypical (attention-deficit hyperactivity disorder and cerebral palsy) development [57,58]. Studying deficits in the flow of multifractality fluctuations longitudinally in typical- and atypical-development might provide insights into the chaotic basis of deficits in perceptual capabilities. Orthotic devices designed to accentuate the flow of fluctuations from distal to proximal body parts could help prevent falls in aging populations. Much like Priplata et al.’s [59,60] successful attempt at supporting posture in the elderly with fractally fluctuating vibrotactile stimulation to the foot sole in contact with the ground, the flow of multifractality fluctuations across the body could be altered to enhance coordination in suprapostural activities. Finally, building distal fluctuations into the architecture of perceptuomotor systems could foster adaptive, flexible chaotic control of robots [61] with dynamic touch capabilities. Our work thus begins to open what could be a broader research program in haptic perception and performance.

